# Human *CEBPA*-N AML exhibits enhanced engraftment and a C/EBP*α*-p30-driven leukemic stem cell program

**DOI:** 10.64898/2026.08.03.742558

**Authors:** Philomina Sona Peramangalam, Mayuresh Konde, Onur Karakaslar, Sebastian Wolf, Shikan Zheng, Akmal Salimov, Sridevi Surapally, Marieke Griffioen, Tongjun Gu, Sridhar Rao, Daniel G Tenen, Thomas Oellerich, Erik van den Akker, Martin Carroll, Caner Saygin, John A. Pulikkan

## Abstract

Leukemic stem cells (LSCs) play a central role in disease progression, therapeutic resistance, and relapse in acute myeloid leukemia (AML). However, the identification and characterization of LSCs remain challenging because of their low abundance and their close phenotypic resemblance to normal hematopoietic stem and progenitor cells. Although patient-derived xenograft (PDX) models have provided important insights into AML biology and LSC heterogeneity, the relative engraftment potential of distinct *CEBPA* mutation subtypes and the immunophenotypic identity of LSCs in *CEBPA* N-terminal mutant AML (*CEBPA*-*N*-AML) remain poorly defined. To address these questions, we compared the engraftment characteristics of primary human CEBPA-mutated AML samples representing the major mutational subtypes using the highly permissive NSGS xenograft model. Primary *CEBPA*-N-AML samples exhibited markedly greater engraftment efficiency and leukemogenic potential than other *CEBPA*-mutated AML subtypes. Furthermore, we identified a CD366⁺CD73⁺CD123⁺CD117⁺CD371⁺CD247⁺ cell population that is highly enriched for functional LSCs in *CEBPA-N*-AML, demonstrating enhanced clonogenic activity, leukemia-initiating capacity, and long-term self-renewal. Collectively, our findings demonstrate that the leukemogenic potential of *CEBPA*-mutated AML is strongly influenced by mutation subtype, with *CEBPA-N*-AML exhibiting superior leukemia-propagating capacity in vivo. We further define a novel immunophenotypic LSC signature specific to *CEBPA-N*-AML, providing new insights into LSC heterogeneity in *CEBPA*-mutated AML and establishing a foundation for the development of LSC-directed therapeutic strategies.

## Introduction

The transcription factor C/EBPα is a master regulator of granulopoiesis. Mice with conditional deletion of *Cebpa* (C/ebpa^Δ/Δ^) lack granulocytes, with a specific block at the common myeloid progenitor (CMP) to granulocyte-monocyte progenitor (GMP) step, highlighting the central role of C/EBPα in granulopoiesis.^1^ The *CEBPA* can translate into two major forms, 42 kD C/EBPα-p42 and 30 kD C/EBPα-p30, based on alternate translation initiation codons.^2^ Mutations in *CEBPA* occur in approximately 10–15% of patients with acute myeloid leukemia (AML).^3–5^ These alterations typically consist of N-terminal frameshift mutations that generate a truncated 30-kDa isoform (C/EBPα-p30) and/or C-terminal mutations affecting the basic leucine zipper (bZIP) domain. Based on mutation status, *CEBPA*-mutated AML can be classified into three molecular subtypes: monoallelic N-terminal mutant (*CEBPA-N*-AML), monoallelic C-terminal mutant (*CEBPA-C*-AML), and biallelic N- and C-terminal mutant *(CEBPA-NC*-AML). Mutation subtype is a major determinant of clinical outcome, with patients harboring *CEBPA-C*-AML or *CEBPA-NC*-AML exhibiting overall survival rates of approximately 60%, compared with only ∼20% for those with *CEBPA-N*-AML.^4^

Although N-terminal *Cebpa* mutations are sufficient to induce AML in genetically engineered mouse models, important biological differences exist between murine and human CEBPA-mutant leukemia. For example, heterozygous *CEBPA-N* mutations cause AML in humans^3,4^ but the corresponding heterozygous mutations do not induce AML in mouse models.^6^ In addition, *Cebpa*-mutant mouse models exhibit significant non-leukemic mortality, raising concerns about their suitability for modeling human disease.^6^ Furthermore, *CEBPA-N* AML occurs predominantly in older adults^7^ and is characterized by substantial interpatient heterogeneity in cooperating mutations, ^4^ clonal architecture,^8^ and leukemia stem cell (LSC) composition, features that are difficult to recapitulate in genetically engineered mouse models. Therefore, patient-derived xenograft (PDX) models provide a more clinically relevant platform for investigating the clonal heterogeneity and LSC hierarchy of human *CEBPA*-mutant AML.

Leukemic stem cells (LSCs) are responsible for disease propagation, therapeutic resistance, and relapse in AML.^9^ However, the identity of LSCs in *CEBPA-*mutant AML and the mechanisms governing their maintenance remain poorly understood. Given that C/EBPα-p30 functions as the primary oncogenic driver in *CEBPA-N*-AML, we hypothesized that its downstream transcriptional program establishes the LSC compartment and that cell-surface genes directly regulated by C/EBPα-p30 could serve as markers of biologically relevant LSCs. Using primary human AML samples and patient-derived xenograft models, we identified a CD366⁺CD73⁺CD123⁺CD117⁺CD371⁺CD247⁺ cell population that is highly enriched for functional LSCs in CEBPA-N-AML, exhibiting enhanced clonogenic activity, leukemia-initiating capacity, and long-term self-renewal.

## Materials and Methods

### Primary human AML samples

Primary human AML samples, including peripheral blood and bone marrow specimens, were obtained from patients treated at the University of Pennsylvania and the University of Chicago.

All patients provided written informed consent in accordance with Institutional Review Board (IRB)-approved research protocols at the respective institutions.

### Mice

All animal experiments were performed in accordance with a protocol reviewed and approved by the Medical College of Wisconsin Institutional Animal Care and Use Committee. NSGS [NOD/SCID-IL2RG–SGM3] mice (Jackson lab, #013062) have been previously described^10^ and maintained at the Medical College of Wisconsin animal facility.

### Colony Forming Unit Assay

Human primary AML cells were resuspended in MethoCult Express (STEMCELL Technologies, #04437), plated at 3,000 cells/mL on a 12-well plate, and cultured for 8 days to form colonies. Colonies were analyzed using an EVOS M7000 microscope.

### Transplantation of human primary AML cells in NSGS mice

Human primary AML cells were T-cell depleted using CD3 Microbeads (Miltenyi Biotec). 1 × 10^6 cells were transplanted into sub-lethally (280 rads) irradiated six- to eight-week-old immunodeficient NSGS mice by tail vein injection. Successful engraftment of edited AML cells was evaluated by flow cytometry of bone marrow aspirates seven days later. The leukemic burden was analyzed by flow cytometry of peripheral blood and bone marrow aspirates five weeks later. Mice were sacrificed after visible characteristics of AML, including reduced motility and grooming activity, hunched back, and pale paws (anemia). For secondary transplantation experiments, 1 × 10^6 cells were transplanted into sub-lethally (280 rads) irradiated six- to eight-week-old immunodeficient NSGS mice by tail vein injection. For LSC transplantation experiments, the number of cells transplanted per human primary AML sample ranged from 12,000 to 70,000. An equal number of LSC+/ LSC-were transplanted into sub-lethally (280 rads) irradiated six- to eight-week-old immunodeficient NSGS mice by tail vein injection

### Flow Cytometry

For flow cytometry, cells were washed twice with 1% BSA in PBS, stained for 20-30 min at 4 °C in the dark, and analyzed using a BD Celesta flow cytometer. For LSC transplantation experiments LSC+ (CD33+ CD366⁺ CD73⁺ CD123⁺ CD117⁺ CD371⁺ CD247⁺) and LSC- (CD33⁺CD366⁻CD73⁻CD123⁻CD117⁻CD371⁻CD247⁻) cells were sorted using BD Aria cell sorter. Flow cytometry analysis was performed using FlowJo Software. The following antibodies were used: Human CD45: Biolegend, #304016, Human CD33: Biolegend, #303413, Human CD366: BD Biosciences, #565566, Human CD247: BD biosciences, #566651, Human CD73: Biolegend, #344003, Human CD371: Biolegend, #353607, Human CD117: Biolegend, ##313227, Human CD123: Biolegend, #30604.

### Mouse histology and tissue sample preparation

Paraffin sections of femur, liver, and spleen were prepared and stained with hematoxylin & eosin for the analysis of tissue architecture. Slides were analyzed using a Nikon Ti2 Widefield inverted microscope, with Nikon 10x, 40x and 100x oil lenses, DS-Ri2; 16.25 MP color camera and NIS-Elements software.

### tSNE Clustering

tSNE Clustering was conducted as reported before.^11^ Briefly, consensus clustering on batch-adjusted gene expression data. A weighted nearest-neighbor graph was constructed using the 2,000 most variable genes (MVGs), which were selected based on median absolute deviation from samples with >70% blast content to minimize the influence of the tumor microenvironment. Cluster assignments were generated using the Leiden algorithm with varying random seeds and resolution parameters, producing 300 clustering iterations for each target cluster number (10–20), yielding a total of 3,300 clustering solutions. A consensus matrix was then generated by calculating the frequency of pairwise sample co-clustering across all 3,300 iterations, resulting in values ranging from 0 to 1. This consensus matrix was converted into a distance matrix (1 − consensus), and hierarchical clustering was performed using Ward’s minimum variance method (Ward.D2).

### Gene expression analysis

Cohort 1^11^ : Quantified gene expression data for the BEAT, TARGET, and TCGA cohorts were obtained from the GDC Data Portal (release 36). To ensure consistent processing across datasets, the Leucegene and LUMC cohorts were reprocessed using the same analysis pipeline. Briefly, FASTQ files were aligned to the GRCh38 reference genome and quantified using STAR with GENCODE v36 gene annotations (60,600 genes). Raw gene count data were batch-corrected using ComBat-Seq, adjusting for cohort, sex, and tissue. Because Leucegene samples were generated using two different library preparation protocols, they were treated as separate batches (Leucegene_stranded and Leucegene_unstranded) during batch correction. Corrected count data were normalized using the geometric mean followed by variance-stabilizing transformation (VST). Residual cohort-specific batch effects were assessed using kBET.

Cohort 2^12^: Bulk RNA-seq generation, alignment, and quantification for the AML discovery cohort have been described^12^. In brief, reads passing FastQC v0.11.9 quality control were aligned with STAR v2.6.1 to the Ensembl GRCh38 (release 82) reference and quantified against Gencode v31 annotations. Count matrices were size-factor normalized (DESeq2 median of ratios) and variance-stabilizing transformed for downstream visualization. Plots were generated using ggplot2 in R v4.5.2, and significance was assessed using the non-parametric Kruskal-Wallis test.

## Results

### Human primary *CEBPA-N* AML cells exhibit superior engraftment and leukemogenic potential in NSGS mice

Multiple studies have investigated engraftment characteristics of multiple AML subtypes using various xenotransplantation models, including the NOD/SCID-IL2RG–SGM3 [NSGS] model.^8,13–17^ Whether *CEBPA* mutant AML differ in their ability to engraft and propagate leukemia in vivo has not been systematically investigated. To determine whether *CEBPA*-mutant AML exhibits distinct engraftment characteristics, we evaluated the engraftment of primary CEBPA-mutant AML samples in NSGS mice. Based on the location of the mutation, samples were classified as *CEBPA-N* (N-terminal mutation), *CEBPA-C* (C-terminal basic leucine zipper [bZIP] mutation), or *CEBPA-NC* (combined N-terminal and C-terminal mutations) (Figure 1A). AML samples harboring mutations in the middle region of *CEBPA (CEBPA-M*) were not included in this analysis. A total of 15 primary human *CEBPA*-mutant AML samples representing the three major *CEBPA* mutational subclasses—*CEBPA-N, CEBPA-C*, and *CEBPA-NC*—were used, with five primary samples included for each subclass.(Figure 1B). T-cell-depleted primary AML cells were transplanted into sublethally irradiated (280 cGy) NOD/SCID-IL2RG–SGM3 [NSGS] recipient mice (Figure 1C), and engraftment was assessed by measuring the percentage of human leukemic blasts in the mouse bone marrow. Remarkably, all *CEBPA-N* AML samples engrafted successfully (25/25), with >0.5% human blasts detected in the bone marrow. In contrast, only one *CEBPA-C* sample and one *CEBPA-NC* sample achieved significant engraftment (Figure 1D). These findings demonstrate that *CEBPA-N* AML possesses a markedly greater engraftment capacity. NSGS mice transplanted with one primary *CEBPA-C* AML sample (5/5) and one primary *CEBPA-NC* AML sample (3/5), both harboring a *WT1* mutation, developed AML (data not shown). All NSGS mice transplanted with *CEBPA-N* AML (25/25) developed AML. Representative flow cytometry plots showing hCD45⁺CD33⁺ human leukemia cells in the bone marrow (BM) and spleen (SP) of moribund NSGS mice transplanted with primary human *CEBPA*-mutant AML cells are shown in Figure 2A. Analysis of mice euthanized at the humane endpoint demonstrated that all *CEBPA-N*-AML-transplanted mice exhibited robust leukemia burden in both the bone marrow and spleen (Figure 2B). Histopathological analyses of the bone marrow, peripheral blood, spleen, and liver confirmed extensive leukemic infiltration in *CEBPA-N*-AML-transplanted mice (Figure 2C). *CEBPA-N*-AML-transplanted mice developed AML with median disease latencies ranging from 20 to 70 days across individual patient samples (Figure 2D). Taken together, these findings establish the robust engraftment and leukemogenic potential of *CEBPA-N* AML in NSGS mice, providing the foundation for subsequent studies investigating its long-term leukemia-propagating and leukemic stem cell properties.

**Figure 1:**
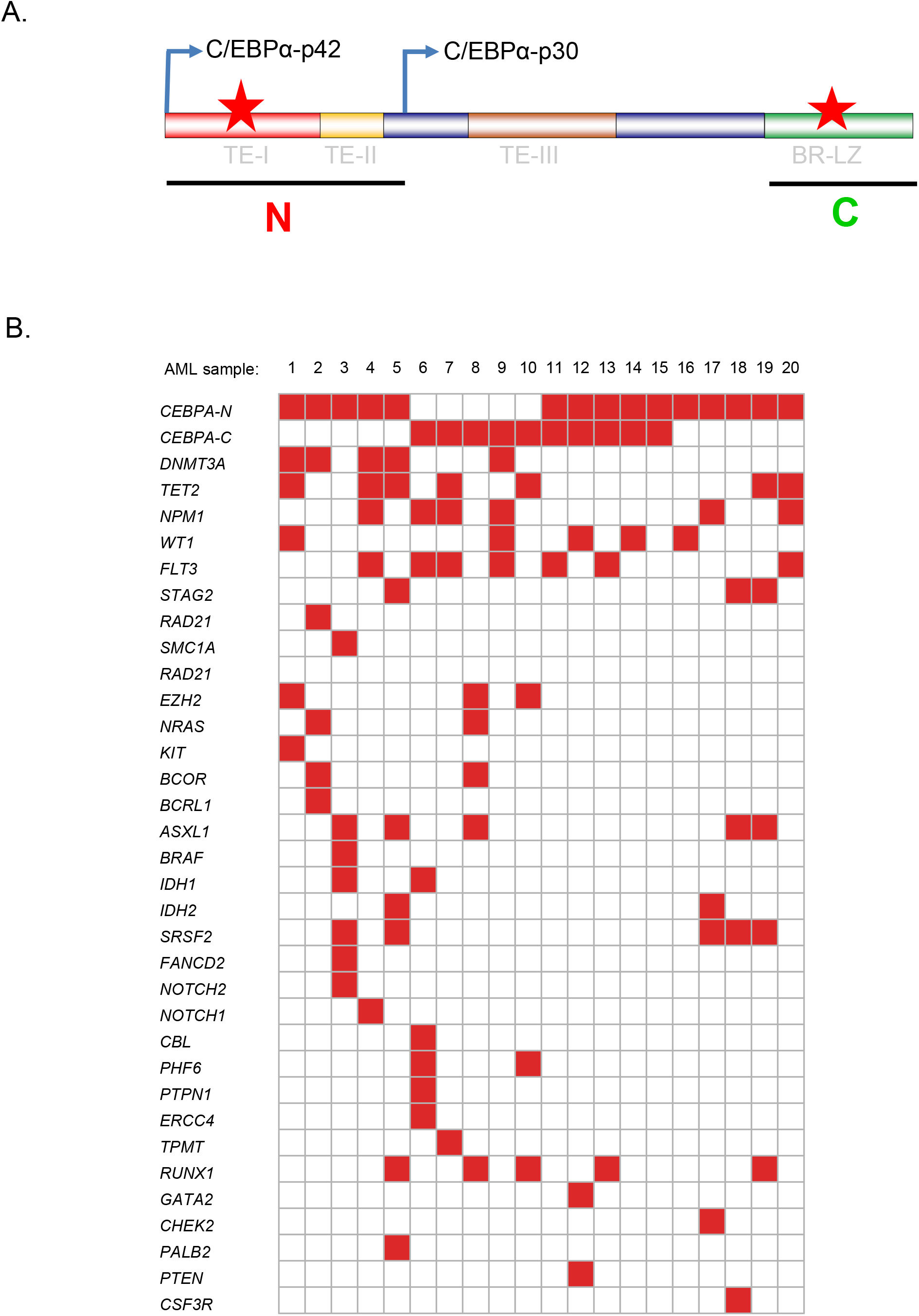
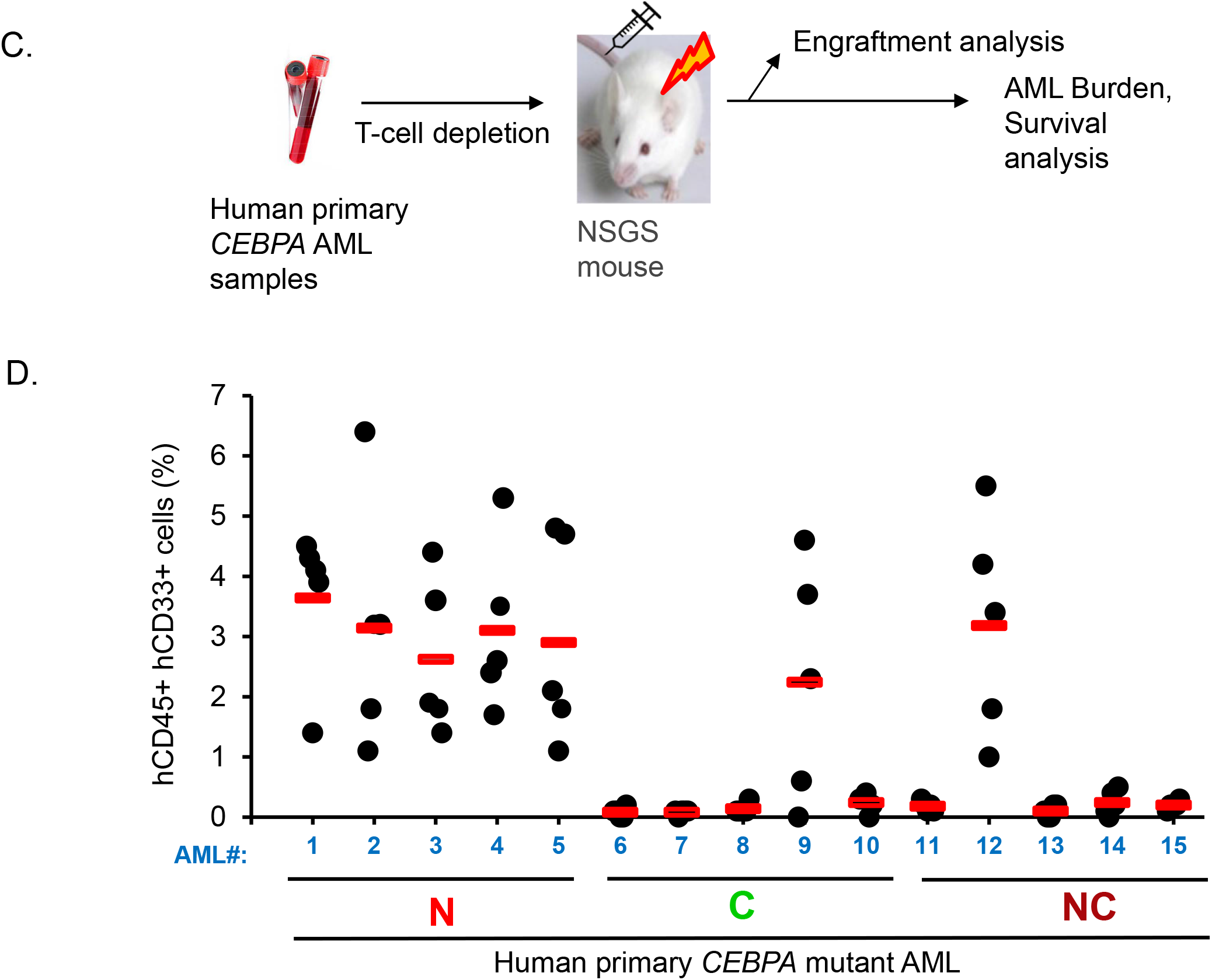
Human primary *CEBPA*-N AML cells exhibit superior engraftment compared with other *CEBPA*-mutant AML subclasses in the NSGS mouse model. (A). Schematic representation of the proteins encoded by wild-type and mutant *CEBPA* alleles. The three transactivation elements (TE-I, TE-II, and TE-III) and the basic region–leucine zipper (bZIP) DNA-binding domain are indicated. The in-frame translation initiation codons at amino acids 1 and 120 generate the full-length C/EBPα-p42 and the truncated C/EBPα-p30 isoforms, respectively. Red stars indicate the locations of the most common *CEBPA* mutations, including N-terminal frameshift mutations that preferentially increase C/EBPα-p30 expression and C-terminal mutations affecting the bZIP domain. (B). Oncoprint illustrating the different types of *CEBPA* mutations and co-occurring genetic alterations in the human primary AML samples included in this study. (C). Schematic representation of the experimental design for transplantation of primary human *CEBPA*-mutant AML cells into NSGS mice. (D). Flow cytometric quantification of hCD45⁺hCD33⁺ cells in NSGS mice bone marrow aspirates 7 days after transplantation with primary human *CEBPA*-mutant AML cells. Each symbol represents an individual mouse, and the red line indicates the group mean.

**Figure 2:**
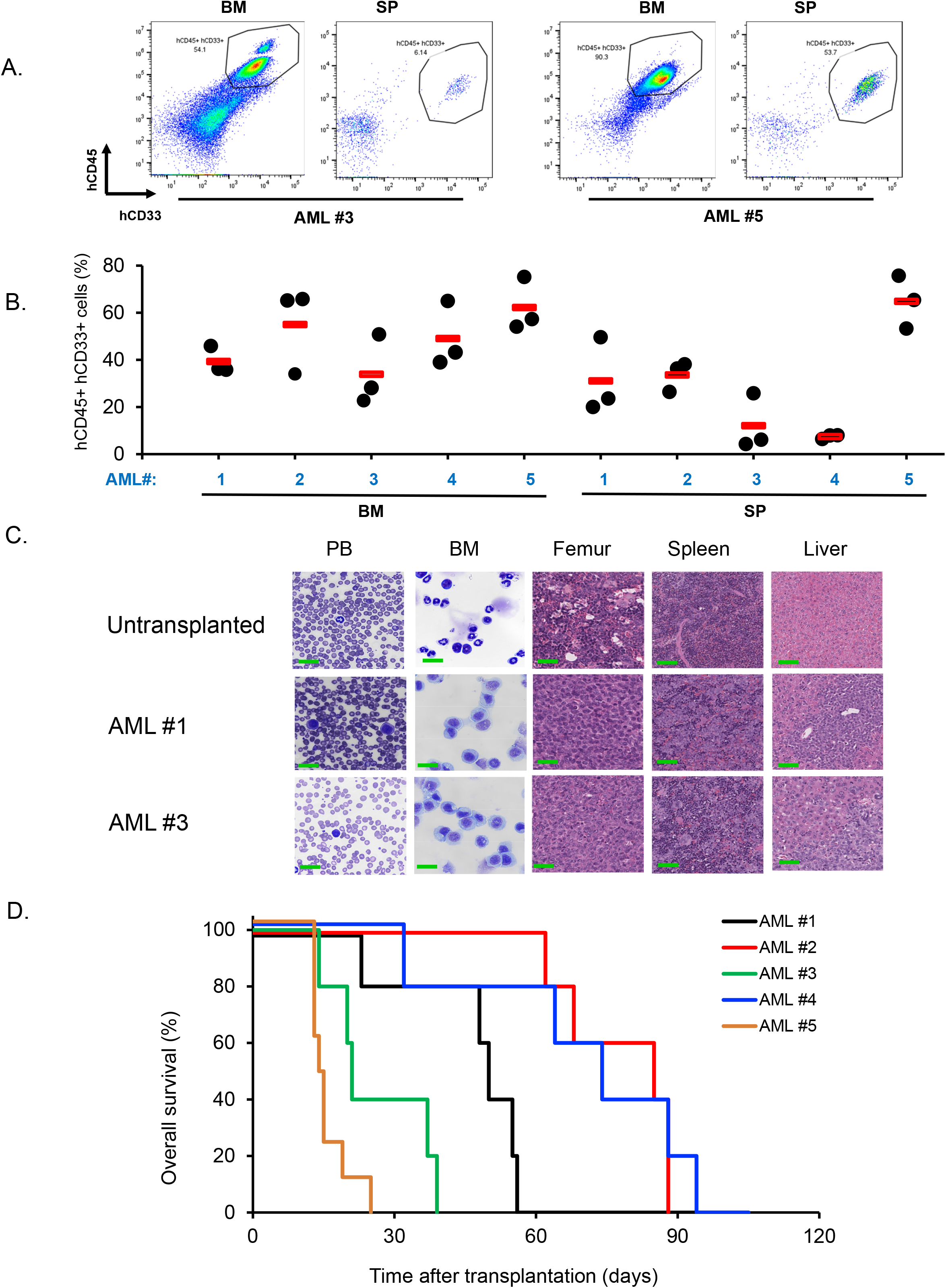
Human primary *CEBPA-N*-AML cells develop AML in the NSGS mouse model. (A). Representative flow cytometry plots showing hCD45⁺CD33⁺ human leukemia cells in the bone marrow (BM) and spleen (SP) of moribund NSGS mice transplanted with primary human *CEBPA*-mutant AML cells. (B). Leukemia burden in moribund NSGS mice engrafted with primary human *CEBPA*-N AML cells, assessed by flow cytometric quantification of hCD45⁺CD33⁺ human leukemia cells in the bone marrow (BM) and spleen (SP). Each symbol represents an individual mouse, and the red line indicates the group mean. (C). Morphological analysis of myeloid blast cells in the peripheral blood (PB) and bone marrow (BM) of untransplanted and moribund NSGS mice transplanted with primary human *CEBPA*-N AML cells, as assessed by May–Grünwald–Giemsa staining. Scale bar, 10 μm. Representative hematoxylin and eosin (H&E)-stained sections of femur, spleen, and liver from untransplanted and moribund *CEBPA*-N AML-transplanted NSGS mice demonstrate disrupted bone marrow and splenic architecture and leukemic infiltration of the liver in transplanted mice. Scale bars: femur, 50 μm; spleen and liver, 100 μm. (D). Kaplan–Meier survival curve of NSGS mice transplanted with primary human *CEBPA*-N AML cells.

### Human *CEBPA-N* AML cells exhibit long-term engraftment capacity in NSGS mice

A defining property of leukemic stem cells is their ability to serially propagate leukemia through secondary transplantation. To evaluate the long-term self-renewal and leukemia-initiating potential of human primary *CEBPA-N* AML cells, leukemic cells harvested from primary NSGS recipients were transplanted into secondary NSGS mice (Figure 3A). We evaluated leukemic engraftment, disease development, and overall survival in the secondary recipients. Analysis of bone marrow from recipient mice euthanized at the humane endpoint demonstrated that all *CEBPA-N*-AML-transplanted mice (15/15) exhibited robust leukemic engraftment (Figure 3B). Histopathological analyses of the bone marrow, peripheral blood, spleen, and liver confirmed extensive leukemic infiltration in the secondary recipient mice, consistent with the findings in primary recipient mice (data not shown). *CEBPA-N*-AML transplanted secondary recipient mice developed AML with median disease latencies ranging from 30 to 80 days across individual samples (Figure 3C). Collectively, these findings demonstrate that human primary *CEBPA-N* AML cells retain the capacity to serially propagate leukemia in NSGS mice, faithfully recapitulating the disease across serial transplantation. These results provide functional evidence for the presence of long-term leukemia-propagating (leukemic stem) cells in CEBPA-N AML.

**Figure 3:**
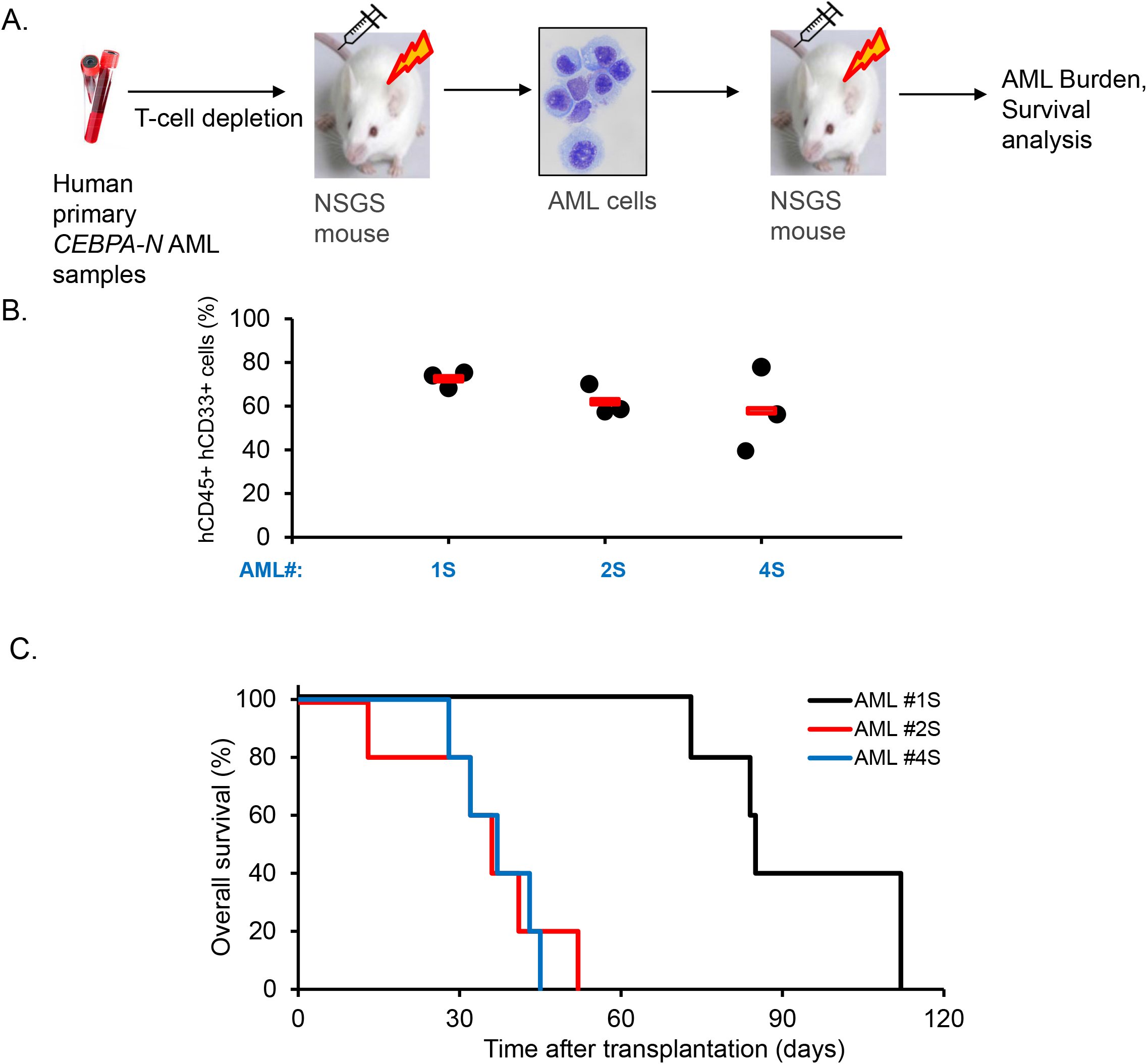
Human *CEBPA*-*N* AML cells exhibit long-term engraftment capacity in NSGS mice. (A). Schematic of the experimental design for secondary transplantation of NSGS-derived mouse *CEBPA*-N AML cells into secondary NSGS recipient mice. (B). Leukemia burden in moribund secondary NSGS recipient mice, assessed by flow cytometric quantification of hCD45⁺CD33⁺ human leukemia cells in the bone marrow. Each symbol represents an individual mouse, and the red line indicates the group mean. (C). Kaplan–Meier survival curve of secondary NSGS recipient mice

### Unsupervised t-SNE Clustering Reveals Unique Molecular Features of CEBPA mutant AML

Given the enhanced leukemogenic potential of *CEBPA-N* AML, we next asked whether this subtype exhibits a distinct global transcriptional profile. To address this question, we analyzed *CEBPA-*mutant AML samples using a previously established t-distributed stochastic neighbor embedding (t-SNE) framework generated from a large multi-cohort AML transcriptomic dataset (n = 1,224).^11^ This approach, developed to identify transcriptionally distinct AML subgroups, provides an unbiased assessment of similarities and differences among AML samples. Previous analyses demonstrated that several AML subtypes, including *CEBPA-C* AML and *CEBPA-NC* AML, form distinct transcriptomic clusters. However, whether *CEBPA-N* AML possesses a unique transcriptional identity and segregates into a distinct cluster has not been investigated. We first generated t-SNE plots encompassing all major AML subtypes together with CEBPA-mutant cases (Figure 4A). We then examined the clustering patterns of the different CEBPA-mutant subclasses. In addition to *CEBPA-N*, *CEBPA-C*, and *CEBPA-NC* AML, this analysis included *CEBPA-M* AML, defined by mutations in the central region of *CEBPA* corresponding to amino acids 121–278. We found that a subset of *CEBPA-N* AML samples clustered with *CEBPA-NC* AML and a subset of *CEBPA-C* AML samples. The remaining *CEBPA-N* AML and *CEBPA-C* AML samples were dispersed across multiple transcriptomic clusters, most prominently within the NPM1-mutant clusters. (Figure 4B). Whether this heterogeneous distribution reflects cooperation between *CEBPA-N* mutations and distinct leukemogenic driver mutations or molecular backgrounds warrants further investigation. Collectively, these findings indicate that *CEBPA-N* AML does not represent a transcriptionally homogeneous AML subtype with a distinct global gene expression profile. Instead, the heterogeneous distribution of *CEBPA-N* AML samples across multiple transcriptomic clusters suggests substantial biological diversity and raises the possibility that *CEBPA-N* mutations cooperate with different leukemogenic driver mutations. These observations prompted us to investigate whether a shared leukemic stem cell program, rather than global transcriptomic features, underlies the aggressive biology of *CEBPA-N* AML.

**Figure 4:**
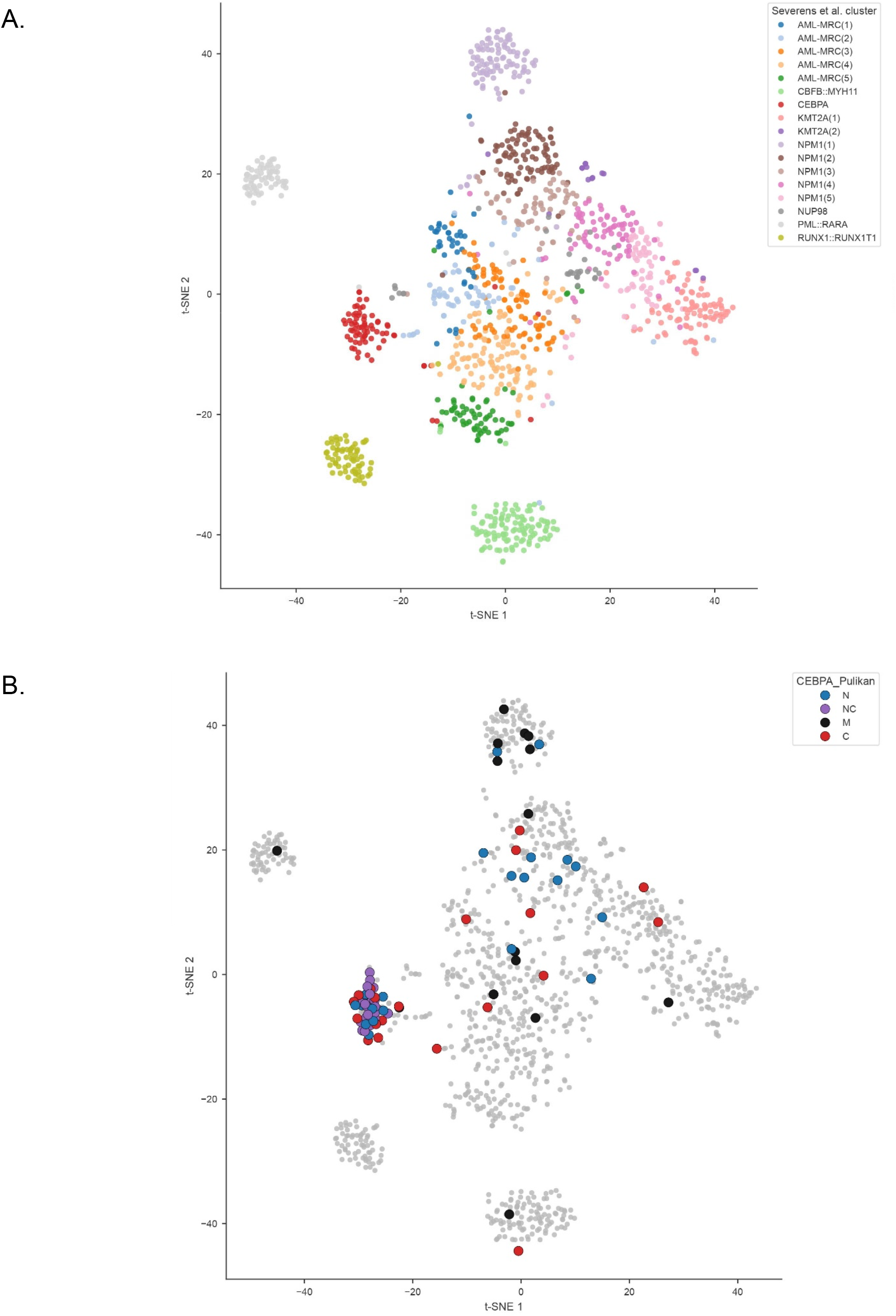
Unsupervised t-SNE Clustering Reveals unique molecular features of *CEBPA* mutant AML. (A-B) t-SNE visualization of gene expression profiles from AML patient samples. Each dot represents an individual AML patient sample. Samples are colored according to AML subtype (A) or *CEBPA* mutation subclass (B).

### Identification of a candidate leukemia stem cell Immunophenotype in *CEBPA-N* AML

Leukemic stem cells (LSCs) are thought to drive AML initiation, propagation, therapeutic resistance, and relapse, making their identification critical for understanding AML biology. Although stemness-associated gene signatures, such as the previously reported 17-gene LSC score,^18^ have provided important insights into LSC biology across multiple AML subtypes, the leukemic stem cell phenotype specific to *CEBPA-N* AML remains undefined. Given the transcriptional heterogeneity observed in *CEBPA-N* AML, we hypothesized that this subtype may instead share a common LSC program. Building on recent studies demonstrating that AML LSCs comprise transcriptionally and epigenetically distinct cellular states^17^, we next sought to identify and characterize the LSC population in *CEBPA-N* AML. To identify candidate LSC markers in *CEBPA-N* AML, we integrated previously reported AML LSC markers^18^ with immunophenotypic markers associated with *CEBPA*-mutant AML identified across multiple independent studies.^19–23^ We further prioritized candidate markers that were direct transcriptional targets of C/EBPα-p30, based on previously published chromatin immunoprecipitation analysis of leukemic cells from a C/EBPα-p30 knock-in mouse model.^19^ This integrative strategy identified the CD366⁺CD73⁺CD123⁺CD117⁺CD371⁺CD247⁺ immunophenotype as a candidate LSC population in CEBPA-N AML (Figure 5A,B). Analysis of individual marker expression within the CD33⁺ blast population of primary human *CEBPA-N* AML samples revealed variable expression frequencies for each marker (Figure 5C). In contrast, cells co-expressing all six markers comprised only 0.05–0.9% of the CD33⁺ blast population, consistent with the expected rarity of leukemic stem cells (Figure 5D). Representative flow cytometric analyses of the six markers in CD33⁺ and CD33⁻ AML blast populations are shown in Figure 5E. Collectively, these findings identify a rare CD366⁺CD73⁺CD123⁺CD117⁺CD371⁺CD247⁺ cell population within human *CEBPA-N* AML with an immunophenotype consistent with leukemic stem cells. By integrating established AML LSC markers with C/EBPα-p30-regulated target genes, we established a rational strategy for identifying the candidate LSC compartment in *CEBPA-N* AML, which was subsequently evaluated by functional assays.

**Figure 5:**
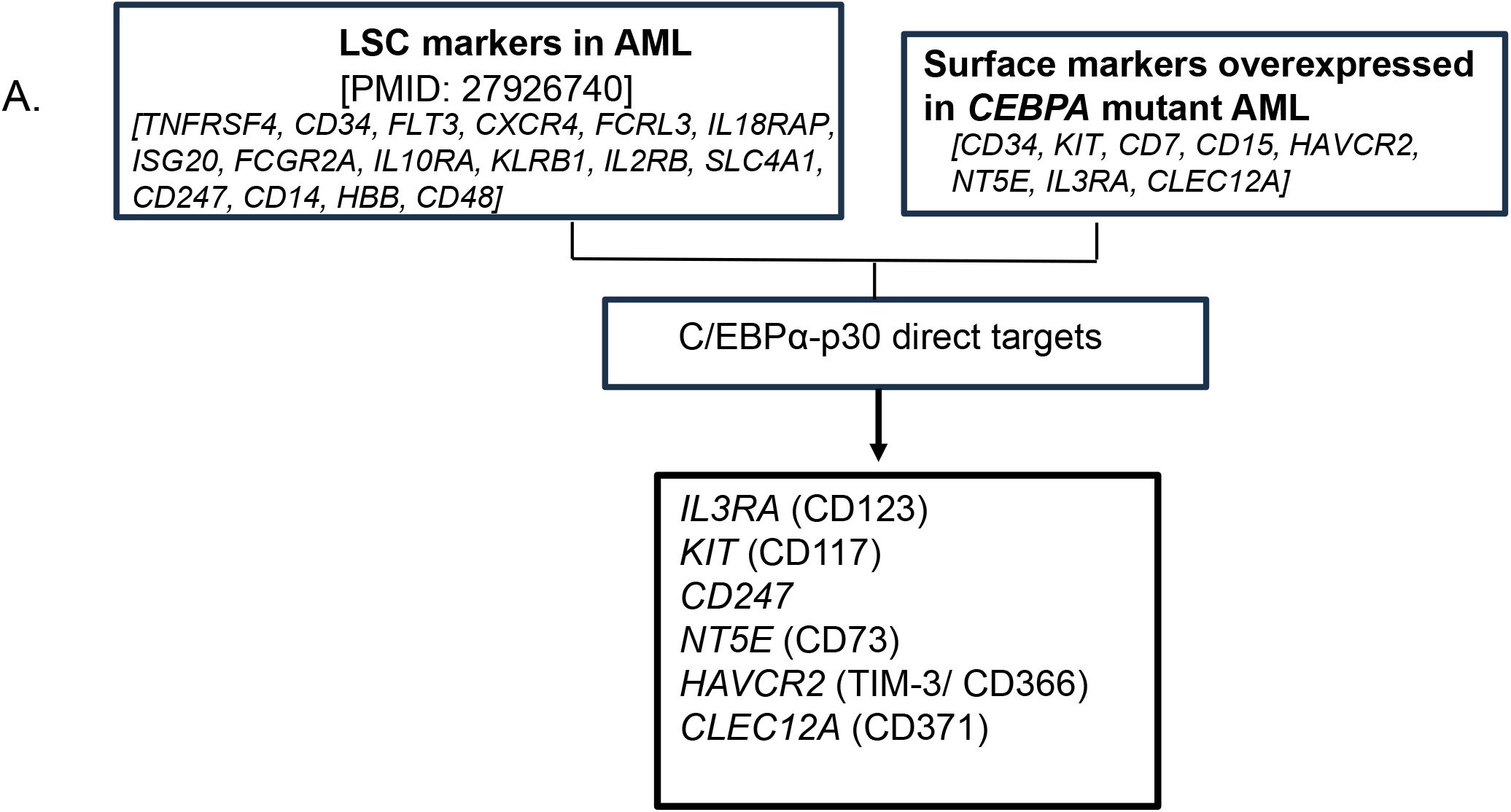
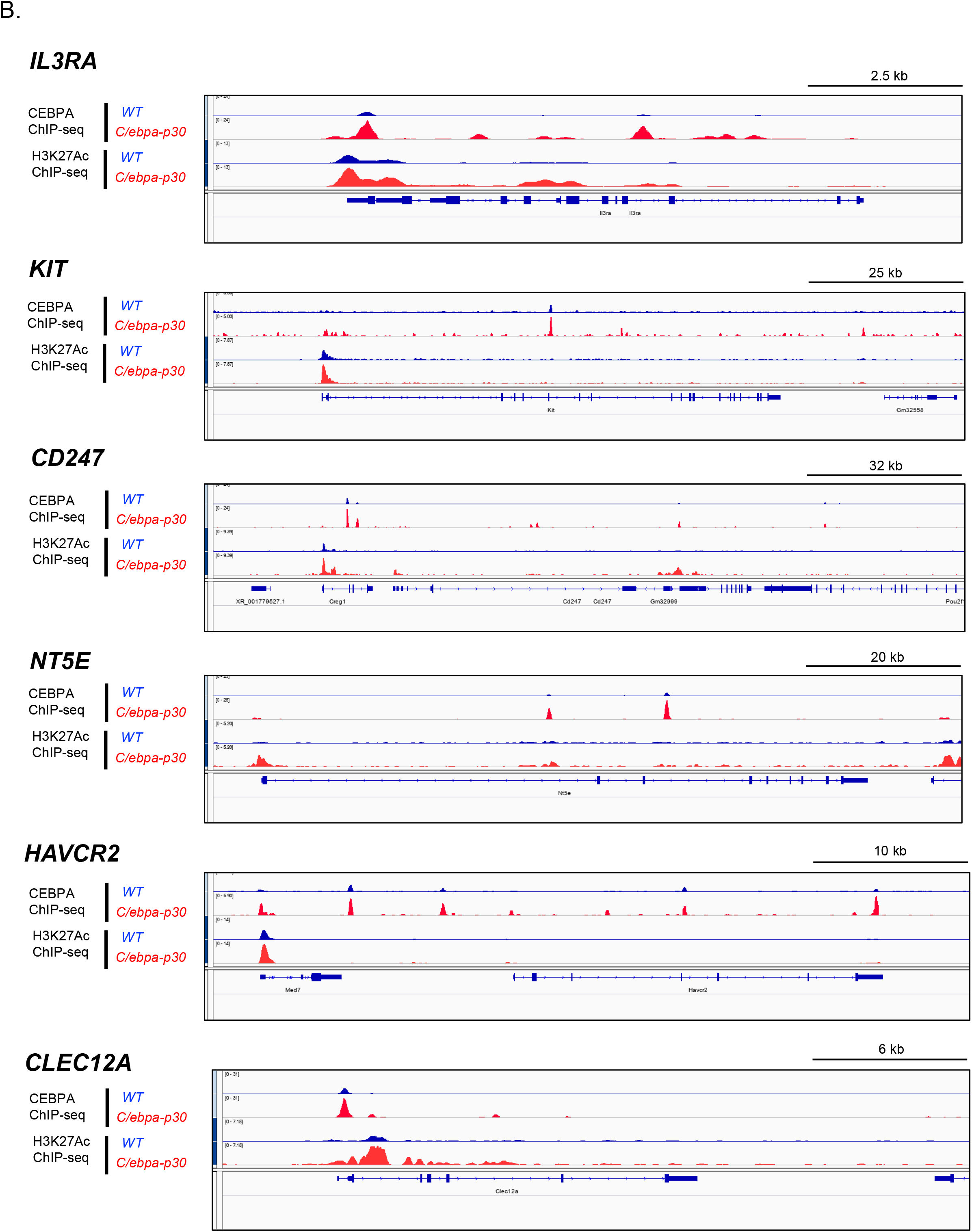
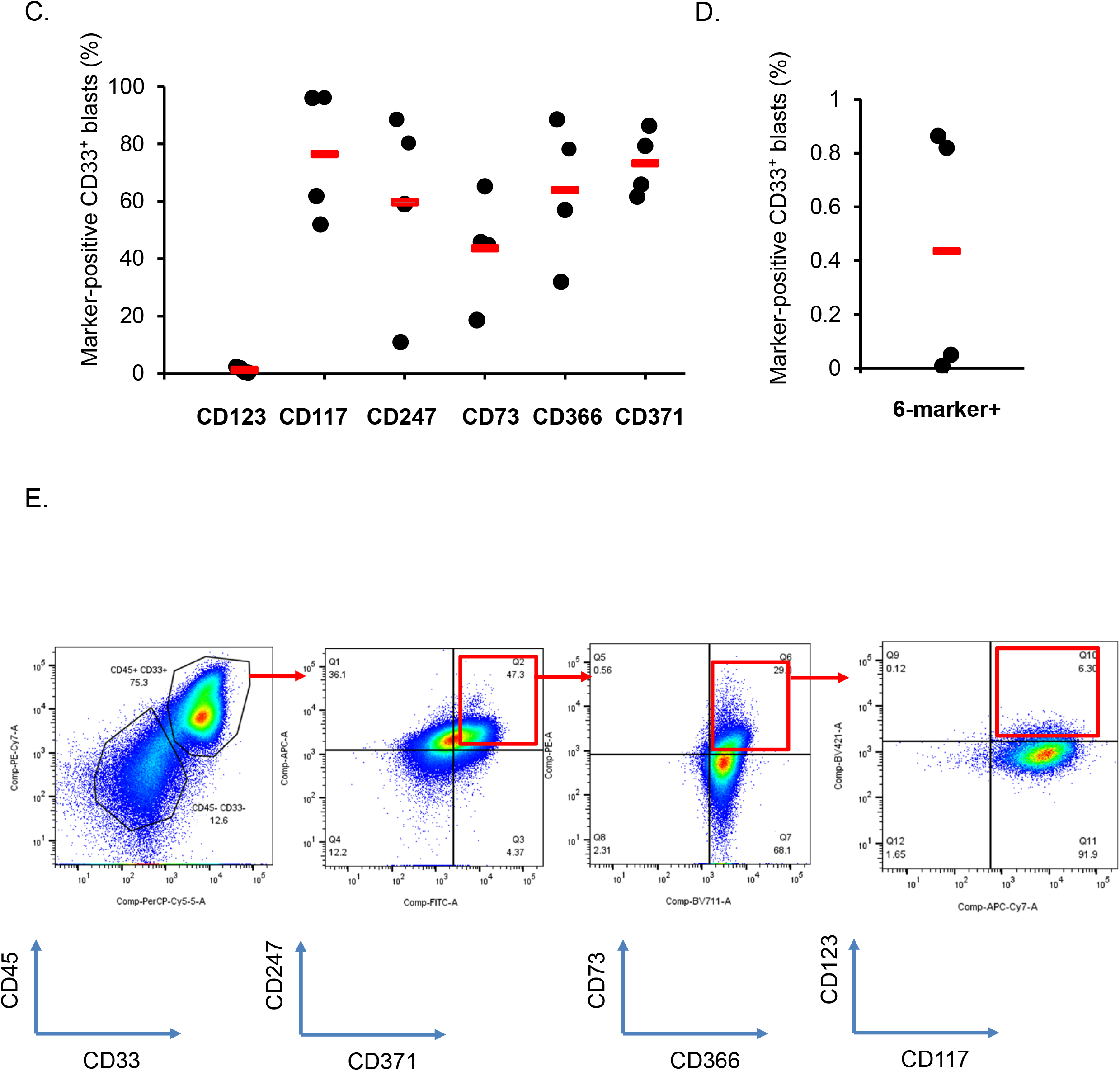
Identification of a candidate leukemia stem cell immunophenotype in CEBPA-N AML. (A). Previously reported AML leukemic stem cell (LSC) markers were integrated with immunophenotypic markers associated with *CEBPA*-mutant AML identified across multiple independent studies. Candidate markers were further prioritized based on direct transcriptional regulation by C/EBPα-p30, leading to the identification of a CD366⁺CD73⁺CD123⁺CD117⁺CD371⁺CD247⁺ immunophenotype as a candidate LSC population in *CEBPA*-N AML. (B). Representative Integrative Genomics Viewer (IGV) tracks showing C/EBPα ChIP-seq and H3K27ac ChIP-seq profiles in granulocyte–macrophage progenitor (GMP) cells from wild-type (WT)and *Cebpa*-p30 leukemic (p30) mice. The tracks demonstrate C/EBPα-p30 occupancy and H3K27ac enrichment at the regulatory regions of candidate LSC marker genes. (C). Flow cytometric quantification of the individual expression of CD366, CD73, CD123, CD117, CD371, and CD247 within the CD33⁺ cell population of primary human *CEBPA*-N AML samples. Each dot represents an individual patient sample, and the red line indicates the group mean. (D). Flow cytometric quantification of the CD366⁺CD73⁺CD123⁺CD117⁺CD371⁺CD247⁺ cell population within the CD33⁺ fraction of primary human *CEBPA*-N AML samples. Each dot represents an individual AML patient sample, and the red line indicates the group mean. (E). Representative flow cytometry plots showing the CD366⁺CD73⁺CD123⁺CD117⁺CD371⁺CD247⁺ cell population within the CD33⁺ fraction of primary human *CEBPA*-N AML samples.

### Candidate leukemic stem cell markers are not selectively enriched in bulk *CEBPA*-mutant AML

Having identified a candidate LSC immunophenotype in *CEBPA-N* AML, we next asked whether the corresponding genes are preferentially expressed in *CEBPA*-mutant AML compared with other AML subtypes. To address this question, we analyzed the expression of these candidate LSC markers in AML samples with a normal karyotype lacking *CEBPA* mutations and compared them with *CEBPA*-mutant AML using two independent transcriptomic cohorts. With the exception of *KIT*, none of the candidate LSC markers exhibited significantly different expression between normal-karyotype AML samples with or without *CEBPA* mutations (Figure 6). In contrast, *KIT* expression was significantly higher in *CEBPA-N, CEBPA-C*, and *CEBPA-NC* AML samples than in normal-karyotype AML lacking *CEBPA* mutations. The absence of differential expression for the remaining candidate markers may have several explanations. First, these genes may represent LSC-associated programs that are shared across multiple normal-karyotype AML subtypes rather than being unique to *CEBPA*-mutant AML. Second, the transcriptomic datasets analyzed were generated from bulk AML samples rather than purified LSC populations, potentially masking subtype-specific differences in gene expression within the rare LSC compartment. Collectively, these findings indicate that the candidate *CEBPA-N* AML LSC markers are not broadly overexpressed at the bulk transcriptome level in *CEBPA-*mutant AML relative to other normal-karyotype AMLs, with the exception of KIT. These results suggest that the biological significance of these markers is more likely related to their expression within the rare LSC compartment than to their overall abundance in bulk leukemic cells. This observation further underscores the importance of functionally validating these candidate markers in purified cell populations.

**Figure 6:**
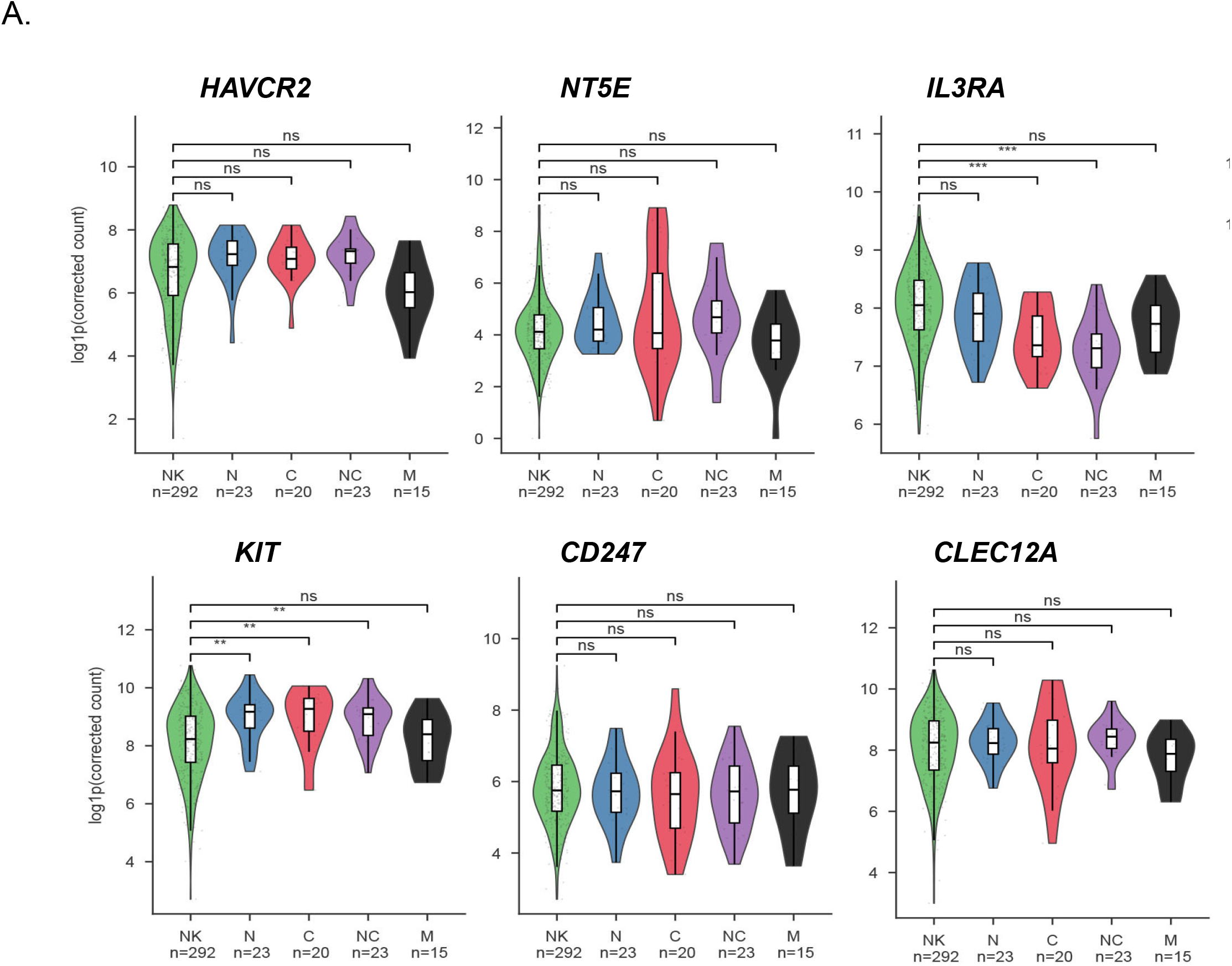
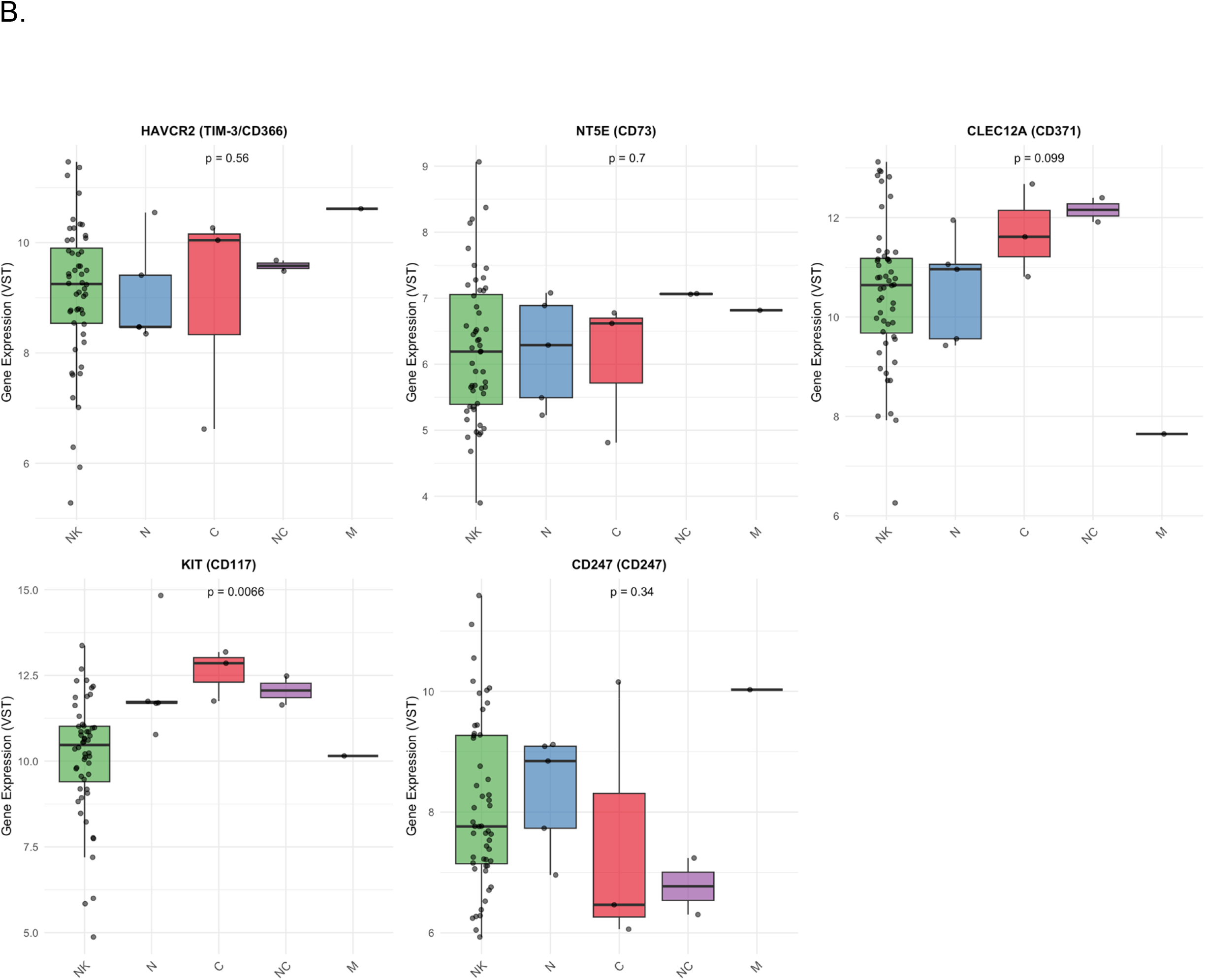
Candidate leukemic stem cell markers are not selectively enriched in bulk *CEBPA*-mutant AML. Normalized mRNA expression levels of *HAVCR2*, *NT5E*, *IL3RA*, *KIT*, *CD247*, and *CLEC12A* in normal-karyotype AML (NK) lacking *CEBPA* mutations and in the indicated *CEBPA* mutation subclasses cohort 1 (A) and cohort 2 (B).

### CD366*⁺*CD73*⁺*CD123*⁺*CD117*⁺*CD371*⁺*CD247*⁺* cells define functional leukemic stem cells in *CEBPA-N* AML

Having identified a candidate LSC immunophenotype in *CEBPA-N* AML, we next investigated whether cells expressing these markers exhibit the defining functional properties of leukemic stem cells. Because C/EBPα-p30 directly regulates multiple cell-surface genes, we hypothesized that coordinated expression of these targets, rather than expression of any single marker, would more accurately identify the leukemia stem cell compartment. To this end, CD33⁺ blast cells co-expressing CD366, CD73, CD123, CD117, CD371, and CD247 (designated LSC⁺) or lacking expression of these markers (LSC⁻: CD33⁺CD366⁻CD73⁻CD123⁻CD117⁻CD371⁻CD247⁻) were isolated from primary human CEBPA-N AML samples by flow cytometric sorting and subjected to colony-forming unit (CFU) assays, xenotransplantation studies, and transcriptional profiling (Figure 7A). CFU assays demonstrated that the clonogenic capacity of LSC⁺ cells was approximately 8- to 10-fold greater than that of LSC⁻ cells (Figure 7B,C). Moreover, colonies generated from LSC⁺ cells exhibited a compact morphology, whereas colonies derived from LSC⁻ cells were more diffuse, consistent with enhanced proliferative activity of the LSC⁺ population. To determine whether the CD366⁺CD73⁺CD123⁺CD117⁺CD371⁺CD247⁺ fraction is enriched for leukemia-initiating cells, purified LSC⁺ and LSC⁻ cells were transplanted into NSGS recipient mice. Mice receiving LSC⁺ cells consistently developed AML with disease characteristics comparable to those observed following transplantation of unfractionated primary *CEBPA-N* AML cells (Figure 7D). Furthermore, leukemic cells recovered from primary recipient mice successfully propagated AML following serial transplantation into secondary NSGS recipients, demonstrating durable leukemia-propagating activity (data not shown). In contrast, NSGS mice transplanted with LSC⁻ cells failed to develop leukemia. Collectively, these findings demonstrate that the CD366⁺CD73⁺CD123⁺CD117⁺CD371⁺CD247⁺ cell fraction is highly enriched for functional leukemic stem cells with enhanced clonogenic, leukemia-initiating, and long-term self-renewal capacities. These results provide functional validation of the candidate LSC markers identified in *CEBPA-N* AML.

**Figure 7:**
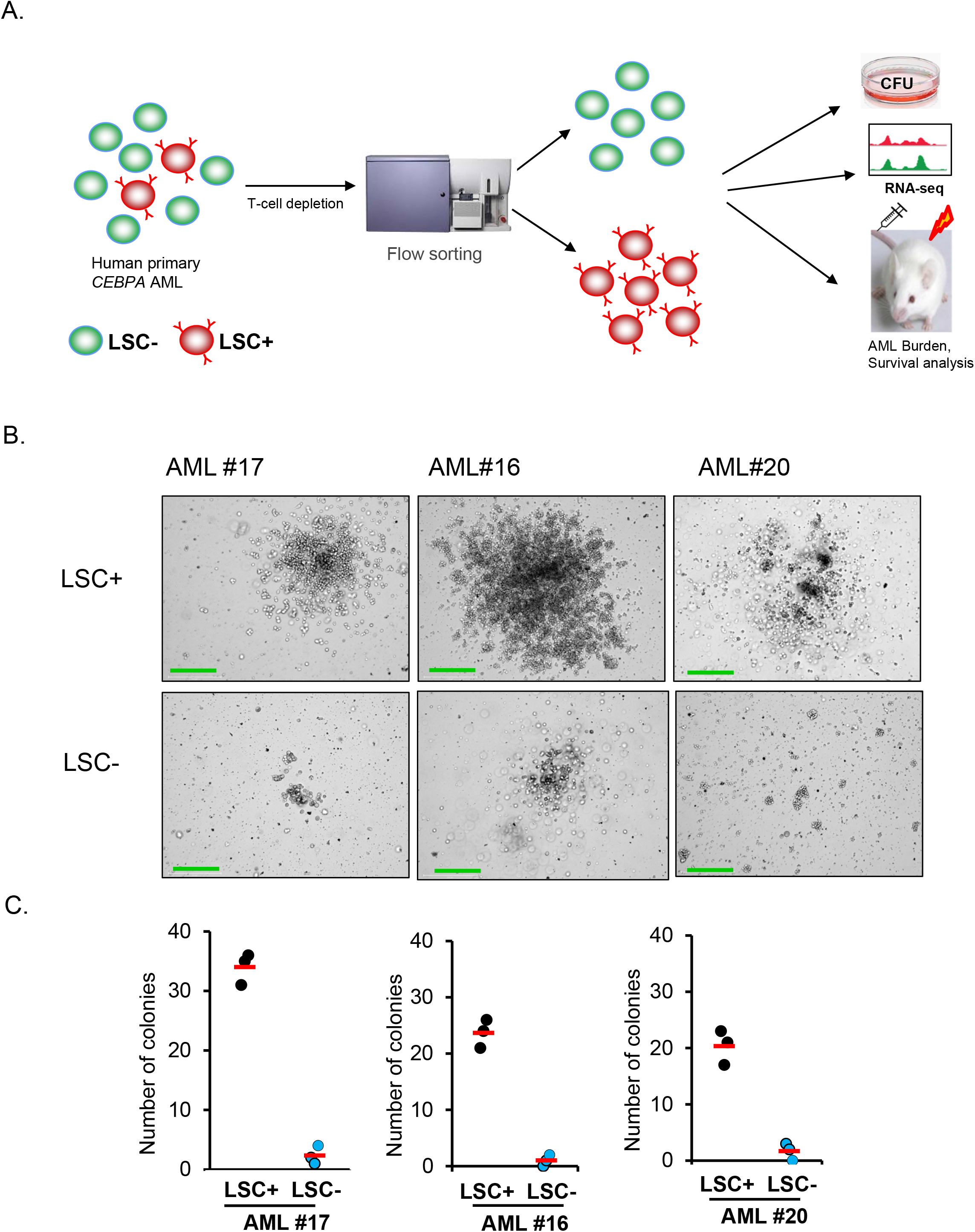
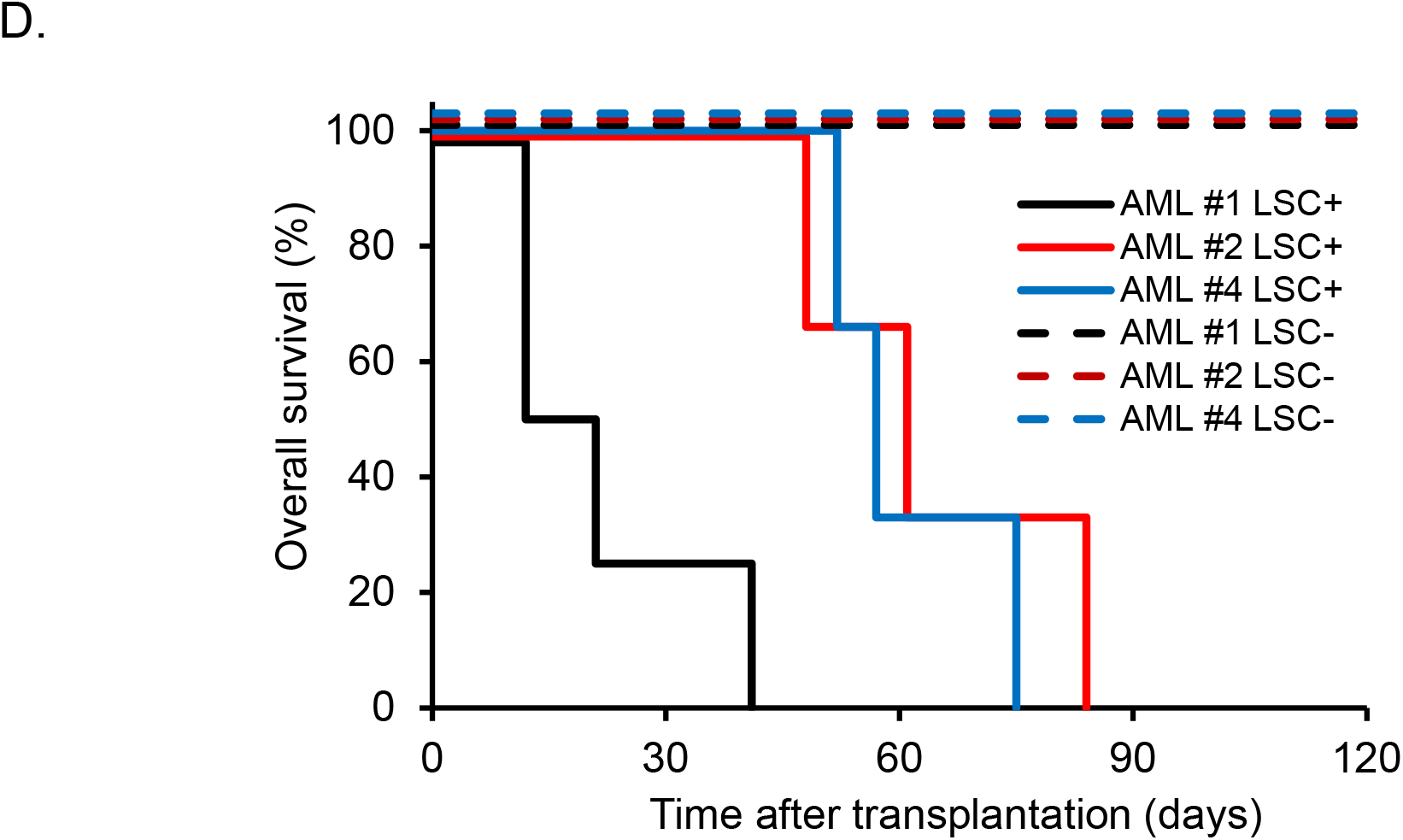
CD366*⁺*CD73*⁺*CD123*⁺*CD117*⁺*CD371*⁺*CD247*⁺* cells define functional leukemic stem cells in *CEBPA*-*N* AML. (A). Schematic illustrating the experimental workflow for functional validation of the candidate LSC markers in *CEBPA*-N AML. (B). Representative images of colonies formed by LSC⁺ [CD33⁺CD366⁺CD73⁺CD123⁺CD117⁺CD371⁺CD247⁺] and LSC⁻ [CD33⁺CD366⁻CD73⁻CD123⁻CD117⁻CD371⁻CD247⁻] cells in methylcellulose colony-forming assays (top). Scale bar, 275 μm. (C). Colony quantification from methylcellulose colony-forming assays performed with LSC⁺ and LSC⁻ cells (bottom). Data are representative of three independent experiments. Each dot represents one replicate, and the red line indicates the group mean. (D). Kaplan–Meier survival curves of NSGS mice transplanted with LSC⁺ or LSC⁻ cells.

## Discussion

The development of innovative therapeutic strategies remains an unmet clinical need in the treatment of AML. Although patient-derived xenograft (PDX) models have provided important insights into AML biology and leukemic stem cell (LSC) heterogeneity in multiple AML subtypes, the relative engraftment potential of the different CEBPA-mutant AML subclasses has remained poorly understood. In this study, we demonstrate that primary *CEBPA-N* AML samples exhibit markedly greater engraftment efficiency and leukemogenic potential than the other *CEBPA-*mutant AML subclasses. These findings establish a robust PDX platform for the preclinical evaluation of novel therapeutic strategies targeting this high-risk AML subtype and provide a valuable model for investigating the biological mechanisms underlying its aggressive clinical behavior.

Our t-SNE analysis showed that a subset of *CEBPA-N* AML samples clustered with *CEBPA-C* and *CEBPA-NC* AML samples, whereas the remaining *CEBPA-N* AML samples were distributed across multiple transcriptomic clusters. This heterogeneity suggests that the transcriptional landscape of *CEBPA-N* AML may depend on whether the *CEBPA N*-terminal mutation represents the initiating (founder) lesion or is acquired as a secondary event during leukemogenesis. Consistent with this hypothesis, several of the non-clustered *CEBPA-N* AML samples co-localized with NPM1-mutant AML, suggesting that cooperating mutations substantially influence the global transcriptional state of these leukemias. Supporting this interpretation, recent genomic studies have reported that more than 40% of *CEBPA-N* AML cases harbor concurrent *NPM1* mutations.^4^ Despite this transcriptomic heterogeneity, all *CEBPA-N* AML samples shared a common functional LSC phenotype. The consistent enrichment of the LSC-associated surface markers and the superior engraftment capacity of LSC-positive cells suggest that C/EBPα-p30 establishes a conserved stemness program that is maintained irrespective of the broader transcriptional background. We therefore propose that C/EBPα-p30 acts as a central regulator of the leukemic stem cell program in *CEBPA-N* AML, whereas cooperating mutations primarily shape the overall transcriptional landscape without disrupting this core stem cell network.

The cancer stem cell model provides the conceptual framework for AML, in which leukemic stem cells (LSCs) constitute a rare self-renewing population responsible for leukemia initiation, maintenance, therapeutic resistance, and relapse. A high LSC burden is associated with poor clinical outcomes and an increased risk of disease recurrence, underscoring the importance of identifying and therapeutically targeting these cells. However, LSC identification remains challenging because of their low abundance, phenotypic and functional heterogeneity, and substantial overlap with normal hematopoietic stem and progenitor cell markers. Although early studies identified the CD34⁺CD38⁻ fraction as the principal LSC compartment^24,25^, subsequent work demonstrated that LSC activity can also reside within CD34⁺CD38⁺ and CD34⁻ populations, highlighting the remarkable diversity of LSC phenotypes across AML subtypes. Additional LSC-associated markers, including CD123, CD93, and GPR56, have also been reported.^26–28^ Despite the well-established role of C/EBPα-p30 in leukemogenesis, the immunophenotypic identity of LSCs in *CEBPA-N* AML has remained poorly defined. Here, we identified a rare CD366⁺CD73⁺CD123⁺CD117⁺CD371⁺CD247⁺ cell population that is highly enriched for functional LSCs, exhibiting enhanced clonogenicity, leukemia-initiating capacity, and long-term self-renewal. These findings establish a distinct C/EBPα-p30-driven LSC population in *CEBPA-N* AML.

Although each C/EBPα-p30-regulated surface marker was expressed in approximately 30–50% of leukemic blasts, simultaneous expression of all six markers was detected in fewer than 1% of cells. This finding is consistent with the hierarchical organization of AML, in which LSCs comprise a rare subpopulation within the bulk leukemic blasts. Similar to the identification of normal hematopoietic stem cells, where combinations of surface markers provide substantially greater specificity than individual markers, our results demonstrate that the coordinated expression of multiple C/EBPα-p30 target genes identifies a highly enriched LSC population in *CEBPA-N* AML. These findings suggest that C/EBPα-p30 does not regulate a single defining LSC marker but instead orchestrates a coordinated surface-marker program that specifies leukemia stem cell identity and function. Although our study focused on cells co-expressing all six markers, it remains possible that smaller marker combinations may also enrich for functional LSCs. Defining the minimal marker set required for optimal LSC isolation will be an important objective for future mechanistic studies and the development of clinically applicable strategies for LSC detection and therapeutic targeting.

Several of the markers identified in this study are themselves attractive therapeutic targets. TIM-3 (CD366) is expressed on AML LSCs but is largely absent from normal hematopoietic stem cells, where TIM-3–galectin-9 signaling promotes LSC self-renewal. Our finding that TIM-3 is part of a C/EBPα-p30-regulated transcriptional program suggests that patients with *CEBPA-N* AML may represent a molecularly defined subgroup particularly amenable to TIM-3-directed therapies, such as sabatolimab. Similarly, CD73 has previously been identified as a direct C/EBPα-p30 target that promotes tumor-protective adenosinergic signaling, further supporting its therapeutic relevance in this disease subtype.^19^ CD123 is among the best-validated therapeutic targets in AML and is currently being explored using antibody-, antibody-drug conjugate-, bispecific antibody-, and cellular immunotherapy-based approaches. Our findings provide additional rationale for evaluating CD123-targeted therapies in *CEBPA-N* AML. Likewise, CD117 (KIT) and CD371 (CLEC12A/CLL-1) are established AML therapeutic targets, and their identification as C/EBPα-p30-regulated genes further supports their contribution to the *CEBPA-N* AML stem cell phenotype. In contrast, CD247 has not previously been implicated in AML biology. Its aberrant expression in the LSC compartment suggests that *CEBPA-N* AML co-opts immune signaling pathways as part of its stemness program, identifying CD247 as a novel biomarker and potential therapeutic target. Consistent with this concept, both CD247 and CD73 are classically associated with immune cell function, further supporting the emerging notion that leukemic cells exploit immune regulatory pathways to promote disease progression and tissue infiltration.

In summary, our study identifies a distinct C/EBPα-p30-driven leukemic stem cell program that underlies the aggressive biology of *CEBPA-N* AML. The identification of a functionally validated six-marker LSC signature, together with the demonstration of superior leukemogenic potential in patient-derived xenograft models, provides a valuable platform for both mechanistic studies and the preclinical evaluation of LSC-directed therapies. Targeting the C/EBPα-p30-regulated stem cell program may therefore represent a promising strategy to improve outcomes for patients with this high-risk AML subtype.

## Acknowledgments

The authors acknowledge Versiti Blood Research Institute Shared Resources Core Facility (RRID: SCR_025503), supported by the Versiti Blood Research Institute Foundation, for services, instrumentation, and specialist support. We also thank the University of Pennsylvania Stem Cell and Xenograft Core (RRID: SCR_010035) for providing human AML samples.

## References

1. Zhang, P., Iwasaki-Arai, J., Iwasaki, H., Fenyus, M.L., Dayaram, T., Owens, B.M., Shigematsu, H., Levantini, E., Huettner, C.S., Lekstrom-Himes, J.A., et al. (2004). Enhancement of hematopoietic stem cell repopulating capacity and self-renewal in the absence of the transcription factor C/EBP alpha. Immunity 21, 853–863. 10.1016/j.immuni.2004.11.006.

2. Lin, F.T., MacDougald, O.A., Diehl, A.M., and Lane, M.D. (1993). A 30-kDa alternative translation product of the CCAAT/enhancer binding protein alpha message: transcriptional activator lacking antimitotic activity. Proceedings of the National Academy of Sciences of the United States of America 90, 9606–9610.

3. Pabst, T., Mueller, B.U., Zhang, P., Radomska, H.S., Narravula, S., Schnittger, S., Behre, G., Hiddemann, W., and Tenen, D.G. (2001). Dominant-negative mutations of CEBPA, encoding CCAAT/enhancer binding protein-alpha (C/EBPalpha), in acute myeloid leukemia. Nature genetics 27, 263–270. 10.1038/85820.

4. Georgi, J.A., Stasik, S., Kramer, M., Meggendorfer, M., Rollig, C., Haferlach, T., Valk, P., Linch, D., Herold, T., Duployez, N., et al. (2024). Prognostic impact of CEBPA mutational subgroups in adult AML. Leukemia 38, 281–290. 10.1038/s41375-024-02140-x.

5. Pulikkan, J.A., Tenen, D.G., and Behre, G. (2017). C/EBPalpha deregulation as a paradigm for leukemogenesis. Leukemia 31, 2279–2285. 10.1038/leu.2017.229.

6. Kirstetter, P., Schuster, M.B., Bereshchenko, O., Moore, S., Dvinge, H., Kurz, E., Theilgaard-Monch, K., Mansson, R., Pedersen, T.A., Pabst, T., et al. (2008). Modeling of C/EBPalpha mutant acute myeloid leukemia reveals a common expression signature of committed myeloid leukemia-initiating cells. Cancer cell 13, 299–310. 10.1016/j.ccr.2008.02.008.

7. Taube, F., Georgi, J.A., Kramer, M., Stasik, S., Middeke, J.M., Rollig, C., Krug, U., Kramer, A., Scholl, S., Hochhaus, A., et al. (2022). CEBPA mutations in 4708 patients with acute myeloid leukemia: differential impact of bZIP and TAD mutations on outcome. Blood 139, 87–103. 10.1182/blood.2020009680.

8. Kawashima, N., Ishikawa, Y., Kim, J.H., Ushijima, Y., Akashi, A., Yamaguchi, Y., Hattori, H., Nakashima, M., Ikeno, S., Kihara, R., et al. (2022). Comparison of clonal architecture between primary and immunodeficient mouse-engrafted acute myeloid leukemia cells. Nature communications 13, 1624. 10.1038/s41467-022-29304-6.

9. Thomas, D., and Majeti, R. (2017). Biology and relevance of human acute myeloid leukemia stem cells. Blood 129, 1577–1585. 10.1182/blood-2016-10-696054.

10. Wunderlich, M., Chou, F.S., Sexton, C., Presicce, P., Chougnet, C.A., Aliberti, J., and Mulloy, J.C. (2018). Improved multilineage human hematopoietic reconstitution and function in NSGS mice. PloS one 13, e0209034. 10.1371/journal.pone.0209034.

11. Severens, J.F., Karakaslar, E.O., van der Reijden, B.A., Sanchez-Lopez, E., van den Berg, R.R., Halkes, C.J.M., van Balen, P., Veelken, H., Reinders, M.J.T., Griffioen, M., and van den Akker, E.B. (2024). Mapping AML heterogeneity - multi-cohort transcriptomic analysis identifies novel clusters and divergent ex-vivo drug responses. Leukemia 38, 751–761. 10.1038/s41375-024-02137-6.

12. Jayavelu, A.K., Wolf, S., Buettner, F., Alexe, G., Haupl, B., Comoglio, F., Schneider, C., Doebele, C., Fuhrmann, D.C., Wagner, S., et al. (2022). The proteogenomic subtypes of acute myeloid leukemia. Cancer cell 40, 301–317 e312. 10.1016/j.ccell.2022.02.006.

13. Krevvata, M., Shan, X., Zhou, C., Dos Santos, C., Habineza Ndikuyeze, G., Secreto, A., Glover, J., Trotman, W., Brake-Silla, G., Nunez-Cruz, S., et al. (2018). Cytokines increase engraftment of human acute myeloid leukemia cells in immunocompromised mice but not engraftment of human myelodysplastic syndrome cells. Haematologica 103, 959–971. 10.3324/haematol.2017.183202.

14. Da Ros, A., Peloso, A., Longo, G., Benetton, M., Indio, V., Cairo, S., Sandri, M., Buldini, B., Bresolin, S., Rosato, A., et al. (2026). Advancing precision therapy in pediatric acute myeloid leukemia through PDX models and mitochondrial targeting. Blood advances 10, 2153–2167. 10.1182/bloodadvances.2025018002.

15. Diaz de la Guardia, R., Velasco-Hernandez, T., Gutierrez-Aguera, F., Roca-Ho, H., Molina, O., Nombela-Arrieta, C., Bataller, A., Fuster, J.L., Anguita, E., Vives, S., et al. (2021). Engraftment characterization of risk-stratified AML in NSGS mice. Blood advances 5, 4842–4854. 10.1182/bloodadvances.2020003958.

16. Morita, K., Wang, F., Jahn, K., Hu, T., Tanaka, T., Sasaki, Y., Kuipers, J., Loghavi, S., Wang, S.A., Yan, Y., et al. (2020). Clonal evolution of acute myeloid leukemia revealed by high-throughput single-cell genomics. Nature communications 11, 5327. 10.1038/s41467-020-19119-8.

17. Boutzen, H., Murison, A., Oriecuia, A., Bansal, S., Arlidge, C., Wang, J.C.Y., Lupien, M., Kaufmann, K.B., and Dick, J.E. (2024). Identification of leukemia stem cell subsets with distinct transcriptional, epigenetic and functional properties. Leukemia 38, 2090–2101. 10.1038/s41375-024-02358-9.

18. Ng, S.W., Mitchell, A., Kennedy, J.A., Chen, W.C., McLeod, J., Ibrahimova, N., Arruda, A., Popescu, A., Gupta, V., Schimmer, A.D., et al. (2016). A 17-gene stemness score for rapid determination of risk in acute leukaemia. Nature 540, 433–437. 10.1038/nature20598.

19. Jakobsen, J.S., Laursen, L.G., Schuster, M.B., Pundhir, S., Schoof, E., Ge, Y., d’Altri, T., Vitting-Seerup, K., Rapin, N., Gentil, C., et al. (2019). Mutant CEBPA directly drives the expression of the targetable tumor-promoting factor CD73 in AML. Science advances 5, eaaw4304. 10.1126/sciadv.aaw4304.

20. Jan, M., Chao, M.P., Cha, A.C., Alizadeh, A.A., Gentles, A.J., Weissman, I.L., and Majeti, R. (2011). Prospective separation of normal and leukemic stem cells based on differential expression of TIM3, a human acute myeloid leukemia stem cell marker. Proceedings of the National Academy of Sciences of the United States of America 108, 5009–5014. 10.1073/pnas.1100551108.

21. Perna, F., Berman, S.H., Soni, R.K., Mansilla-Soto, J., Eyquem, J., Hamieh, M., Hendrickson, R.C., Brennan, C.W., and Sadelain, M. (2017). Integrating Proteomics and Transcriptomics for Systematic Combinatorial Chimeric Antigen Receptor Therapy of AML. Cancer cell 32, 506–519 e505. 10.1016/j.ccell.2017.09.004.

22. Liu, Q., Qi, L., Yang, M., Zhang, X., Li, F., Wei, H., and Wang, J. (2023). Immunophenotype distinctions of CEBPA mutation subtypes in acute myeloid leukemia. International journal of laboratory hematology 45, 743–750. 10.1111/ijlh.14124.

23. Mannelli, F., Ponziani, V., Bencini, S., Bonetti, M.I., Benelli, M., Cutini, I., Gianfaldoni, G., Scappini, B., Pancani, F., Piccini, M., et al. (2017). CEBPA-double-mutated acute myeloid leukemia displays a unique phenotypic profile: a reliable screening method and insight into biological features. Haematologica 102, 529–540. 10.3324/haematol.2016.151910.

24. Lapidot, T., Sirard, C., Vormoor, J., Murdoch, B., Hoang, T., Caceres-Cortes, J., Minden, M., Paterson, B., Caligiuri, M.A., and Dick, J.E. (1994). A cell initiating human acute myeloid leukaemia after transplantation into SCID mice. Nature 367, 645–648. 10.1038/367645a0.

25. Bonnet, D., and Dick, J.E. (1997). Human acute myeloid leukemia is organized as a hierarchy that originates from a primitive hematopoietic cell. Nature medicine 3, 730–737. 10.1038/nm0797-730.

26. Jin, L., Lee, E.M., Ramshaw, H.S., Busfield, S.J., Peoppl, A.G., Wilkinson, L., Guthridge, M.A., Thomas, D., Barry, E.F., Boyd, A., et al. (2009). Monoclonal antibody-mediated targeting of CD123, IL-3 receptor alpha chain, eliminates human acute myeloid leukemic stem cells. Cell stem cell 5, 31–42. 10.1016/j.stem.2009.04.018.

27. Iwasaki, M., Liedtke, M., Gentles, A.J., and Cleary, M.L. (2015). CD93 Marks a Non-Quiescent Human Leukemia Stem Cell Population and Is Required for Development of MLL-Rearranged Acute Myeloid Leukemia. Cell stem cell 17, 412–421. 10.1016/j.stem.2015.08.008.

28. Pabst, C., Bergeron, A., Lavallee, V.P., Yeh, J., Gendron, P., Norddahl, G.L., Krosl, J., Boivin, I., Deneault, E., Simard, J., et al. (2016). GPR56 identifies primary human acute myeloid leukemia cells with high repopulating potential in vivo. Blood 127, 2018–2027. 10.1182/blood-2015-11-683649.

